# SNPLift: Fast and accurate conversion of genetic variant coordinates across genome assemblies

**DOI:** 10.1101/2023.06.13.544861

**Authors:** Eric Normandeau, Maxime de Ronne, Davoud Torkamaneh

**Affiliations:** Institut de Biologie Intégrative et des Systèmes (IBIS), Université Laval, Québec, Canada; Département de Phytologie, Université Laval, Québec, Canada; Centre de recherche et d’innovation sur les végétaux (CRIV), Université Laval, Québec, Canada; Institut intelligence et données (IID), Université Laval, Québec, Canada

## Abstract

**Motivation:** The advent of high-throughput sequencing technologies and the availability of reference genomes have provided an unprecedented opportunity to discover and genotype millions of genetic variants in hundreds or even thousands of samples. Variant calling, the identification of genetic variants from raw sequencing data, is both time-consuming and computationally demanding. Currently, reference genomes are evolving very rapidly and new assembly versions come out more and more frequently. To take advantage of new or improved reference genomes, raw reads alignments, genotype calling, and filtration must typically all be redone. This is a costly and time consuming operation that is not always viable when projects are under time constraints.

**Results:** Here, we introduce SNPLift, a bioinformatic pipeline that can quickly transfer the coordinate of nucleotide variants (SNPs and Indels) between different versions of reference genomes. We tested SNPLift on nine SNP datasets in VCF format from different species (*Homo sapiens, Arabidopsis thaliana, Coregonus clupeaformis, Medicato truncatula, Oriza sativa, Salvelinus namaycush, Solanum lycopersicum, Zea mays, and Glycine max*). Depending on the species, we achieved accurate lifting of variants ranging from 92.92% to 99.69%. Importantly, SNPLift significantly reduces the computational resources and time required for variant analysis compared to performing a complete re-analysis using a new reference genome. SNPLift offers a fast and efficient solution to leverage the benefits of updated or improved reference genomes.

**Availability and implementation:** SNPLift is available at https://github.com/enormandeau/snplift with its documentation. It contains a script that runs an automated test on a small dataset, composed of 190,443 SNPs in chromosome 1 of *Medicago truncatula*. SNPLift uses only common tools that are easy to install and works under Linux and MacOS.

## Introduction

The availability of reference genomes and the advent of high-throughput sequencing technologies has created an exceptional opportunity to systematically detect genetic variants like single nucleotide polymorphisms (SNPs) and small insertions and deletions (indels) in a wide array of organisms (Eichler, 2019). This has provided crucial knowledge about genomic diversity and architecture and has paved the way for subsequent genetic studies. Traditionally, genetic variation has been identified by comparing individual resequenced genomes to a single reference genome (Torkamaneh, Boyle, *et al*., 2018). However, decreasing costs, enhanced scalability, and increasing quality of sequencing technologies, along with advancements in computational tools and resources, have accelerated rapid generation of both new and improved versions of reference genomes (Kaye and Wasserman, 2021). Currently, there are 9,176 species listed in GenBank with three or more assembled genome versions and, of these, 3,488 have 10 or more distinct versions (National Center for Biotechnology Information [NCBI]; accessed March 2023). The availability of improved or new reference genomes brings about the necessity of re-analyzing genetic variants, which can be a time-consuming and computationally intensive task, particularly when dealing with a large number of sequenced individuals. To overcome this challenge, we present SNPLift, a bioinformatic pipeline designed to efficiently transfer coordinates, from VCF or other file formats, from one version of a genome to another. SNPLift enables the rapid utilisation of the valuable resources provided by updated reference genomes, mitigating the need for extensive and resource-intensive re-analyses.

## Materials and Methods

SNPLift uses python 3.5+ (van Rossum, 1995), R 3+ (R Core Team, 2021), bash 4+(Bash (3.2. 48) - GNU Project - Free Software Foundation, 2007), gnu parallel (Tange, 2018), git (Chacon and Straub, 2014), bwa (Li and Durbin, 2009), samtools (Danecek *et al*., 2021), minimap2 (Li, 2021), and miniasm (Li, 2016) and requires three input files: two genome assemblies (referred to as old genome and new genome) in Fasta format from the same or a closely related species, as well as a file, for example a VCF or a bed file, containing coordinates from the old genome. The configuration file for SNPLift requires the user to provide the name and location of the input files and contains parameters that can be adjusted. These details, along with comprehensive instructions, can be found on SNPLift’s Github page (https://github.com/enormandeau/snplift). During the execution of SNPLift, there is an optional step that involves aligning both genomes with minimap2 and producing a synteny dot plot to help assess the collinearity of the two genomes. SNPLift then indexes the new genome and proceeds to find the corresponding position of each marker in the new genome. This is achieved by extracting flanking sequences around the positions of the marker in the old genome and mapping these regions (by default 601bp) onto the new genome. From the alignment file, SNPLift extracts features to calculate scores for each marker. Markers with high scores are retained, and a subsequent attempt is made to keep those with lower scores if they meet certain criteria. Specifically, a defined number of additional markers (by default 20) on either side of the marker with a lower score are considered. If the scores of these 21 SNPs are also low, the marker is discarded. For the remaining markers, SNPLift computes the Pearson correlation between the positions of the 21 markers on both genomes. Markers with Pearson correlation above 0.99 are retained. Finally, SNPLift generates an output file containing the retained markers and with their positions in the new genome. The pipeline also provides various options to control the creation of VCFs. All analyses performed for this paper were run on a Linux Mint 19.2 server equipped with Intel Xeon 2.2 GHz cores.

### Input files format

SNPLift relies on the names of chromosomes and/or scaffolds found in the input file to extract short sequences from the old genome. Therefore, the names of chromosomes or scaffolds in the input file must match those found in the old gnome file. If the input file is a VCF, it should adhere to the standard VCF format (https://github.com/samtools/hts-specs/blob/master/VCFv4.4.pdf). Additionally, the genome assemblies should be provided as uncompressed Fasta files.

## Results

To assess the performance of SNPLift, we used nine different datasets (Table 1) that varied in terms of genome size (0.12 to 3.3 Gbp) and number of variants (14.78 thousand to 49.51 million). The data used in this study and its description has been made publicly available on the Dryad repository (doi:10.5061/dryad.h9w0vt4nx). As shown in Table 1, the accuracy of variant lifting ranged from 92.92% to 99.69% depending on the species, with the largest dataset requiring about 90 minutes. Computing time, excluding required time to index the reference genome, was very strongly correlated with the number of variants. For datasets with millions of SNPs, once the genome was indexed, and using 20 cores, SNPLift was able to transfer approximately 0.5 to 1 million variants per minute. This estimation is slightly influenced by the size, composition, and quality of the assembly.

**Table 1.** Accuracy of SNPLift-transferred variants on nine different species.

| Species | Assembly Size (Gb) | Number of SNPs | Transfer (%) | Run Time (m) | Index Time (m) | Total Time (m) | MSPM* |
| --- | --- | --- | --- | --- | --- | --- | --- |
| **Human**<br>(*Homo sapiens*) | 3.3 | 10,737,716 | 93.30 | 12.5 | 69.9 | 82.4 | 0.86 |
| **Arabidopsis**<br>(*Arabidopsis thaliana*) | 0.12 | 12,752,919 | 99.95 | 7.7 | 1.9 | 9.6 | 1.65 |
| **Maize**<br>(*Zea mays*) | 2.18 | 49,510,515 | 99.69 | 50.5 | 39.0 | 89.5 | 0.98 |
| **Rice**<br>(*Oryza sativa*) | 0.37 | 28,092,897 | 95.99 | 26.2 | 7.4 | 33.5 | 1.07 |
| **Soybean**<br>(*Glycine max*) | 0.98 | 4,900,192 | 99.19 | 9.8 | 17.0 | 26.8 | 0.50 |
| **Tomato**<br>(*Solanum lycopersicum*) | 0.73 | 495,419 | 99.42 | 1.1 | 13.9 | 15.0 | 0.44 |
| **Barrel Medic**<br>(*Medicago truncatula*) | 0.43 | 39,980 | 98.43 | 0.5 | 8.3 | 8.8 | 0.09 |
| **Whitefish**<br>(*Coregonus clupeaformis*) | 2.75 | 5,621,210 | 92.92 | 21.0 | 53.8 | 74.9 | 0.27 |
| **Coho salmon**<br>(*Salvelinus namaycush*) | 2.35 | 14,779 | 95.70 | 1.5 | 41.9 | 43.4 | 0.01 |

\*MSPM: Million SNPs Per Minute transferred (excluding the time required for genome indexing)

SNPLift demonstrates a high success rate in transferring variants when both genomes are relatively similar, for example in *Arabidopsis*. However, some situations can lead to a lower success rate if the two genome versions are more distant or from different species where the mapping may not be as effective. Additionally, if the genomes contain a high proportion of paralogous regions, repeated sequences, and transposable elements, fewer markers will be transferred. It is important to acknowledge that due to local differences between assemblies, a small fraction of the markers may have positions that differ by a few base pairs compared to the positions obtained by re-aligning the same samples to the new reference genome.

In order to directly assess the accuracy of SNPLift, a specific analysis was conducted. A dataset consisting of 40 soybean full genomes (Torkamaneh, Laroche, *et al*., 2018) were mapped to versions 2 and 4 of the soybean reference genome, resulting after filtration in the identification of more than 2 million variants. Then, SNPLift was used to transfer the variants’ coordinates from version 2 to version 4 of the genome. We compared the coordinates of the reanalysis and the SNPLift-transferred variants, revealing that 99.61% of the variants were successfully transferred. Among these variants, 20,554 (1.029%) were found to be located at different positions within 300bp, while 10,946 SNPs (0.548%) were within 20bp. It is important to note that these cases may require further improvements in order to enhance transfer accuracy.

## Conclusion

Reference genomes provide useful resources for identification of genetic variation across species. However, as new or improved genome references are generated, the need to re-analyze sequencing data to obtain variant coordinates in the updated assemblies arises. In this study, we demonstrated the effectiveness of SNPLift as a useful tool for rapidly and accurately transferring genetic loci coordinates between different assemblies. As reference genome assemblies continue to be produced at an increasing pace, we hope that SNPLift will prove useful to a broad array of users in different research communities.

## Acknowledgment

The authors wish to thank Génome Québec, Genome Canada, the government of Canada, the Ministère de l’Économie et de l’Innovation du Québec, the Canadian Field Crop Research Alliance, Semences Prograin Inc., Sollio Agriculture, Grain Farmers of Ontario, Barley Council of Canada, and Université Laval.

## Funding

This work was funded by Genome Canada [#6548] under Genomic Applications Partnership Program (GAPP).

## Conflict of Interest

none declared.

## Notes

### Competing Interest Statement

The authors have declared no competing interest.

